# PRE-CLINICAL IMMUNE RESPONSE AND SAFETY EVALUATION OF THE PROTEIN SUBUNIT VACCINE NANOCOVAX FOR COVID-19

**DOI:** 10.1101/2021.07.20.453162

**Authors:** Tran Thi Nhu Mai, Bruce May, Ung Trong Thuan, Nguyen Mai Khoi, Nguyen Thi Thuy Trang, Dinh Van Long, Doan Chinh Chung, Tran The Vinh, Khong Hiep, Nguyen Thi Thanh Truc, Hua Hoang Quoc Huy, Nguyen Viet Anh, Ha Tan Phat, Phan Dang Luu, Nguyen Truong An, Bui Thi Ngoc, Tu Tieu My, Nguyen Thi Theo, Le Thi Thuy Hang, Dong Thi Lan, Huynh Trong Hieu, Ho Phien Huong, Le Nguyen Thanh Thao, Truong Cong Thao, Pham Hoang Phi, Y Luong Cong, Nie Lim, Cao Minh Ngoc, Nguyen Duy Khanh, Trinh Thanh Hung, Do Minh Si

## Abstract

The Coronavirus disease-2019 (COVID-19) pandemic caused by the Severe Acute Respiratory Syndrome coronavirus 2 (SARS-CoV-2), has become a dire global health concern. The development of vaccines with high immunogenicity and safety is crucial for control of the global COVID-19 pandemic and prevention of further illness and fatalities. Here, we report development of SARS-CoV-2 vaccine candidate, Nanocovax, based on recombinant protein production of the extracellular (soluble) portion of the S protein of SARS-CoV-2. The results showed that Nanocovax induced high levels of S protein-specific IgG, as well neutralizing antibody in three animal models including Balb/C mice, Syrian hamsters, and non-human primate (Macaca leonina). In addition, the viral challenge study using the hamster model showed that Nanocovax protected the upper respiratory tract from SARS-CoV-2 infection. No adverse effects were induced by Nanocovax in swiss mice (*Musmusculus* var. Albino), Rats (Rattus norvegicus), and New Zealand rabbits. These pre-clinical results indicated that Nanocovax is safe and effective.

## 1. Introduction

The Coronavirus disease-2019 (COVID-19) pandemic caused by the Severe Acute Respiratory Syndrome coronavirus 2 (SARS-CoV-2), has become a dire global health emergency [1]. Since it was first reported in Wuhan, China at the end of 2019, there have been over 165 million cases worldwide, and nearly 3.5 million deaths as of May 2021 [2], with no obvious short term resolution. Like SARS-CoV (79% genomic sequence identity) [3], SARS-CoV-2 utilizes the receptor angiotensin-converting enzyme 2 (ACE-2), which is expressed on numerous cells including lung, heart, kidney, and intestine cells as the entry fusion receptor by its viral spike, a homotrimeric complex of spike (S) proteins [4]. The S protein is a homomeric class I fusion protein with each S monomer containing the N-terminal S1 subunit, which includes the receptor-binding domain (RBD), and the C-terminal S2 subunit, which is anchored to the viral membrane and is required for trimerization of the spike itself and fusion to host membrane [5]. During fusion to host cell membranes, the S protein undergoes extensive conformational changes that cause dissociation of the S1 subunit from the complex and the formation of a stable post-fusion conformation of the S2 subunit [6]. Therefore, the S protein of SARS-CoV-2 plays a vital role in the invasive process.

Potential vaccines against SARS-CoV-2 have therefore focused on the S protein and include mRNA-lipid nanoparticles that encode the S protein, viral vectored DNA-based vaccines (notably recombinant adenoviruses), and subunit vaccines that contain purified S protein [7][8]. One report from the World Health Organization (WHO) estimated that 102 S protein-targeted vaccines are in clinical development and 185 S protein-targeted vaccines are in pre-clinical development (*COVID-19 Vaccine Tracker and Landscape*). The vaccine candidates that have been developed or are under development include recombinant protein vaccines, whole inactivated vaccines, viral vector-based vaccines, and DNA vaccines to prevent virus infection. Currently, three vaccines are approved by Food & Drug Administration (FDA): BNT162b2 is a lipid nanoparticle-formulated, nucleoside-modified mRNA vaccine (Comirnaty^®^; Pfizer/BionNtech) [10], Moderna COVID-19 vaccine is mRNA-based [11], and Janssen (Johnson and Johnson) [12] is based on an adenovirus vector and is used in adults aged 18 years and older. Besides vaccines, oligonucleotides, peptides, interferon, and small-molecule drugs have been suggested to control infection of SARS-CoV-2 [13]. According to FDA during a public health emergency, three anti-SARS-CoV-2 mAb therapies have been approved for the treatment of non-hospitalized patients with mild-to-moderate COVID-19 including bamlanivimab, etesevimab or casirivimab with imdevimab [14].

In the present study, we have developed a Covid-19 subunit vaccine, named Nanocovax, based on recombinant protein technology to produce the extracellular (soluble) portion of the S protein of SARS-CoV-2. In brief, a gene encoding the S protein was constructed with the wild-type sequence of the S protein extracellular domain. The construct was transfected into CHO cells, named CHO-spike cells, with the highest S protein expression selected. The S protein of SARS-CoV-2 was then absorbed into aluminium hydroxide gel adjuvant (Alhydrogel^®^; Croda, Denmark). We describe the pre-clinical studies of Nanocovax vaccine and illustrate its immunogenicity, efficacy, safety in mice, hamsters, non-human primates, rats, and rabbits.

## 2. Materials and methods

### 2.1. Plasmid construction, cell clone and purification

#### 2.1.1. Plasmid construction

The pCNeoMEM vector was used as the expression vector. The pCNeoMEM plasmid containes a G418 resistance gene used as a selection marker, a MoMLV promoter to express the target gene, a human universal chromatin opening element (UCOE), untranslated regions from the Chinese hamster EEF1A1 gene (eEF1A1), and a synthetic matrix attachment sequence (sMAR). The gene encoding the Spike protein of SARS-CoV-2 (UniProt P0DTC2) codon-optimized for expression in CHO cells was synthesized by Genscript (Piscataway, NJ, USA). The optimized DNA fragment was cloned in the expression vector to create pCNeoMEM-S, which was completely sequenced by NextGen technology at the Center for Computational and Integrative Biology, Harvard University.

#### 2.1.2. Cell culture and protein expression

CHO cells (cGMP bank, Thermo Fisher Scientific, Waltham, MA, USA) were propagated and maintained in the animal component free, chemically defined medium, PowerCHO-2, (Lonza, Walkersville, MD, USA) in a 37°C and 5% CO_2_. Suspension cells were routinely subcultured every 2-3 days at a cell concentration of 2×10^5^ cells/mL. Before transfection, the cells were seeded at 1×10^6^ cells/mL into a 6-well plate and cultured for 24 h. For transfection, Lipofectamine LTX Reagent (Thermo Fisher Scientific, Waltham, MA, USA) was used following manufacturer’s instructions. In optimized conditions, total 3 μg of plasmids was used to transfect into 1×10^6^ cells per well in 6-well plates using PowerCHO-2 medium. After 48 h of transfection, the expression of S protein was evaluated by using ELISA. After transient transfection, selection was performed by culturing transfected cells in PowerCHO-2 medium, supplemented with 400 μg/mL of Geneticin G418 (Sigma-Aldrich, St.Louis, MO, USA). The selective medium was replaced every 2-3 days for 3 weeks to obtain stable cell lines. Then, single-cell cloning was initiated by limiting dilution. The clones with higher productivity were selected to create master cell bank (MCB) and working cell bank (WCB). Process development was performed using the high throughput multi-parallel Bioreactor Ambr 15 and Ambr 250 (Sartorius Stedim Biotech, Göttingen, Germany). Protein production at pilot scale for nonclinical trial material was archived in 500L Bioreactor (BIOSTAT STR®500, Sartorius Stedim Biotech, Göttingen, Germany) using a fed-batch process using PowerCHO-2 medium as basal medium, supplement with Feed 3 (Excellgene, Monthey, Switzerland), trace elements and activated sugars (n=3).

#### 2.1.3. Protein purification

The purification of S protein from cell broth was performed using on AKTA Pilot 600R system (GE Healthcare, Little Chalfont, UK). After harvesting, the cell supernatant was clarified by depth-filtration (Merck Millipore, Darmstadt, Germany) to remove cells and purified by consecutive chromatography steps. Firstly, sample was loaded onto Blue Sepharose column (Cytiva, Marlborough, MA, USA) to specifically collect S proteins and then treated at low pH 3.2 - 3.5 for 60-70 min to inactivate virus. Next, sample was flowed through Q membrane, and then Phenyl membrane (Sartorius Stedim Biotech, Göttingen, Germany) to remove host cell DNA, proteins, and endotoxin. Exotic viruses were removed by nano-filtration. Purified S proteins were exchanged to storage buffer and then filtered through 0.22 um filter (Sartorius Stedim Biotech, Göttingen, Germany). The concentration of purified protein was determined by ELISA. The purity of protein was determined by SE-HPLC, residual host cell DNA assay, and residual host cell protein assay.

### 2.2. Spike protein of SARS-CoV-2 analysis

#### 2.2.1. Spike protein identification

##### 2.2.1.1. SDS-PAGE and Western blot assay

Purified proteins were mixed with loading buffer, and loaded onto SDS-PAGE gels. Proteins were electrophoresed for 110 minutes at 80V, 500mA and visualized using Coomassie Brilliant Blue G250 staining (Sigma-Aldrich, St. Louis, MO, USA) or transferred to a nitro cellulose blotting membrane (Sigma-Aldrich, St. Louis, MO, USA) at a constant current of 250 mA, for 90 minutes. Next, the membranes were blocked in 0.5% BSA (Sigma-Aldrich, St. Louis, MO, USA) and incubated with human anti-S1 antibody (Abcam, Cambridge, MA, UK) for 3 hours at room temperature. After washing, the membranes were incubated with horseradish peroxidase (HRP)-linked rabbit anti human IgG antibody (Abcam, Cambridge, MA, UK) for 1 hour at room temperature. Membranes were washed 3 times with PBST and the protein bands were visualized by enhancing TMB substrate solution (Sigma-Aldrich, St. Louis, MO, USA).

##### 2.2.1.2. ELISA assay

The culture supernatant sample or purified S protein was coated on each well of the microtiter plate overnight at 4°C. After blocking the wells with 1% BSA for 1 hour, dilutions of human anti-S1 antibody (Abcam, Cambridge, MA, UK) were added to wells and incubated for 3 hours at room temperature. After washing, horseradish peroxidase (HRP)-conjugated rabbit anti human IgG antibody (Abcam, Cambridge, MA, UK) was added and incubated for 1 hour at room temperature. Bound antibodies were detected using TMB substrate (Sigma-Aldrich, St. Louis, MO, USA). Absorbance was read at 450 nm on Multimode Plate Reader (Promega, Madison, WI, USA)

#### 2.2.2. Structure analysis

##### 2.2.2.1. Molecular mass assay

The molecular mass of purified protein was measured by MALDI-TOF-MS at Biofidus AG Company. In brief, recombinant SARS-COV-2 Spike protein was desalted and concentrated with C4 ZipTips (Merck Millipore, Darmstadt, Germany) and spotted on a ground steel target using 2’,5’-Dihydroxyacetophenone. The sample was measured via MALDI-TOF-MS (UltrafleXtreme, Bruker Daltonik GmbH, Bremen Germany) in positive ion mode. Recorded MS spectra were processed by the software FlexAnalysis (Bruker Daltonik GmbH, Bremen Germany).

##### 2.2.2.2. Peptide mapping assay

Peptide mapping of purified protein was performed by Biofidus AG Company. Briefly, the peptide mapping of a recombinant SARS-CoV-2 spike protein was measured by LC-ESI-MS using different digestion strategies. The focus of the peptide mapping was sequence verification and analysis of N- and C-terminal modifications. The mass spectrometric analysis was performed with a Compact QTOF mass spectrometer (Bruker Daltonik GmbH, Bremen Germany). The recorded LC-ESI-MS and -MS/MS spectra were processed, annotated and searched against a customized sequence database using Mascot (Matrix Science). Modified peptides were identified by their exact mass and retention time and quantified by their mass spectrometric signal intensity.

### 2.3. Formulation of Nanocovax vaccine

Aluminum hydroxide gel adjuvant (alhydrogel^®^ 1.3%) in this study was obtained commercially from Croda (Denmark) (#21645-51-2) and re-autoclaved according to the manufacturer’s instruction. The antigen S protein of SARS-CoV-2 was diluted in saline buffer with 20 mM of phosphate. For antigen formulation, 1 ml of S protein was mixed with aluminum hydroxide gel adjuvant and stirred at 200 RPM for 18 hours at cool temperature (2-8°C). Finally, one dose contained 25 μg; 50 μg; 75 μg; and 100 μg of S protein plus 0.5 mg of Al^3+^.

### 2.4. *In vivo* immunigenicity evaluation of Nanocovax vaccine in Balb/c mice, Syrian hamsters, and non-human primate models

#### 2.4.1. Animal vaccination

Balb/c mice of both sexes, 6-10 weeks old were used for immunological studies. Syrian hamsters (*Mesocricetus auratus*) of both sexes, 8-12 weeks old were used for immunological studies. Other model, northern pig-tailed macaques (*Macaca leonina*), male, 4-5 years old, 7-9 kg were also used for immunological study. They were housed in temperature-controlled rooms at the animal facilities with fixed light-dark cycles (12-hour shifts). Upon arriving, animals were quarantined and let to acclimate to housing conditions for at least one week before being used. The Balb/c mice, Syrian hamsters, as well non-human primates were immunized intramuscularly with Nanocovax at doses of 25 μg, 50 μg, 75 μg, and 100 μg, their serum samples were collected quantification of spike protein specific IgG antibodies by ELISA. Blood samples were collected and let to clot at room temperature for 60 min, then centrifuged at 1000 x g for 15 min. Upper serum fraction was collected and heat-inactivated at 56°C in 30 minutes before use or kept at −20°C.

#### 2.4.2. Quantification of SARS-CoV-2 spike protein specific IgG antibody by ELISA

The method of quantifying SARS-CoV-2 spike protein-specific IgG antibody was adapted from Tan C. W. et al, [15]. Briefly, S protein was diluted in PBS to a final concentration of 1000 ng/mL. The 96 well plate was coated by adding 100 μL of coating buffer (1000 ng/mL of S protein) per well, sealed and incubated at 4°C for overnight or at room temperature for 2h, then washed 3 times with wash buffer. After washing step, each well was blocked with 300 μL of blocking buffer (PBS + 2% BSA) and incubated at room temperature (RT) for 2 hours. After blocking step, wells were washed 3 times with washing buffer and loaded with 100 μL of standard solutions (SARS-CoV-2 Spike S1 subunit antibody (#1035206; R&D Systerms) and serum samples and incubated at RT for 2 hours. Wells were washed 3 times with washing buffer then 100 μL of diluted goat anti-mouse IgG (Fc specific)-Peroxidase antibody (#A0168; Sigma-aldrich) were added. After incubating 1 hour, 100 μL of TMB substrate was added. The plate was kept in dark at RT for 5 minutes for color development. Color development reaction was stopped by adding 100 μL of 1M H_2_SO_4_ solution. The absorbance was measured as the optical density 450 nm.

#### 2.4.3. *In vitro* surrogate virus neutralization assay

The virus neutralization ability of antibodies in sera of Balb/C mice, hamster, and non-human private were determined by the virus surrogate neutralization kit (cat # L00847, Genscript, Singapore). The percent of neutralizing virus in sera were determinded according to the manufacturer’s protocol

#### 2.4.4. *In vitro* SARS-CoV-2 virus neutralization assay (PRNT)

The PRNT assay detects and quantifies neutralizing antibody SARS-CoV-2 in serum samples. Sera were 2-fold serially diluted in culture medium with a starting dilution of 1:20. The diluted sera were mixed with 100 plaque-forming units (PFU) of SARS-CoV-2 virus for 1 h at 37 °C. The virus–serum mixtures were added onto Vero E6 cell monolayers and incubated 1 h at 37°C in 5% CO2 incubator. Then the plates were overlaid with 1% agarose in cell culture medium and incubated for 4 days when the plates were fixed and stained. Antibody titres were defined as the highest serum dilution that resulted in>50% (PRNT50) reduction in the number of plaques. PRNT was performed in duplicate using 24-well tissue culture plates in a biosafety level 3 facility (BSL3) in the National Institute of Hygiene and Epidemiology (NIHE), Hanoi, Vietnam, adapted from Okba N. et al,. [16]

### 2.5. Protective efficacy evaluation of Nanocovax vaccine in Syrian hamsters

#### 2.5.1. Animal vaccination

Syrian hamsters (*Mesocricetus auratus*) of both sexes, 8-12 weeks old were used for the viral challenge study. They were housed in temperature-controlled rooms at the animal facilities with fixed light-dark cycles (12-hour shifts). Upon arriving, animals were quarantined and let to acclimate to housing conditions for at least one week before being used. Syrian hamsters were immunized intramuscularly with two doseages of Nanocovax 50 μg on day 0 and day 7.

#### 2.5.2. Viral Challenge study

All procedures were performed in BSL-3 facility for animals at NIHE (National Institute of Hygiene and Epidemiology) Vietnam. We used the hCoV-19/Vietnam/CM99/2020 isolate which was isolated in NIC-NIHE. The isolate was propagated in Vero E6 to prepare virus stocks. Titers of virus stocks were determined by PRNT_50_

The hamsters were assigned to the following groups: (1) vaccinated with Nanocovax on day 0 and day 7 and then challenged with the high level of the SARS-CoV-2 virus on day 14 by the intranasal route (TCID_50_ = 2×10^5^); (2) vaccinated with Nanocovax on day 0 and day 7 and then challenged with the low level of the SARS-CoV-2 virus on day 14 by the intranasal route (TCID_50_ = 1×10^3^), and (3) placebo injection with PBS and challenged with the high/low level of the SARS-CoV-2 virus on day 14 by the intranasal route (TCID_50_ = 2×10^5^ and 1×10^3^). Baseline body weights were measured before infection. Animals were monitored for signs of morbidity (such as weight loss, ruffled hair, sweating, etc.) for 14 days. On day 28, their lungs were collected for detection of SARS-CoV-2 by Real-time RT-PCR. Quantitative detection of SARS-CoV-2 on the lung samples was adapted from WHO’s protocol [17]. Infection doses were chosen to base on Imai et al,. [18].

#### 2.5.3. Real-time RT-PCR

In our research, qRT-PCR was performed to quantify SARS-CoV-2. This PCR amplifies the *envelope* (E) gene of SARS-CoV-2 using forward primer 5’-ACAGGTACGTTAATA-GTTAATAGC-3’; reverse primer: 5’-ATATTGCAGCAGTACGCACAC-3’; and probe: 5’-FAM-ACACTAGCCATCCTTACTGCGCTTCG-BBQ-3’. The Real-time RT-PCR assays were conducted using TagMan One-Step RT-PCR kit (Thermo Fisher Scientific, Waltham, MA, USA) on Real-Time PCR System (Biorad, CA, USA) with the following cycling conditions: 55°C for 10 min for reverse transcription, 95°C for 3 min, and 45 cycles of 95°C for 15 s and 58°C for 30 s. The absolute copy number of viral loads was determined using serially diluted DNA control targeting the E gene of SARS-CoV-2.

#### 2.5.4. Histological examination

Tissue samples from lung of hamster were processed overnight at room temperature for formalin fixation and dehydration following a established protocol. The fixed specimens were embedded directly in paraffin blocks and cut into sections with 3–5 μm thickness using a rotary microtome (Leica RM 2125RTS, Leica Biosystems Nussloch GmbH, Nussloch, Germany) and then stained with HE solution. The mounted specimens were observed and photomicrographs were taken using an optical microscope with DP27 digital camera (Olympus, Tokyo, Japan). The histopathologic examination was performed by a pathologist.

### 2.6. Safety evaluation of Nanocovax vaccine in rodent animals

#### 2.6.1. Single-dose toxicity test

According to ICH/GLP guidelines with minor modification, single-dose toxicity study was conducted on adult male and female mice with minor modifications. The mice were fasted overnight but with free access to water prior to treatment. A total of 60 mice of both sexes were divided into 06 groups (n = 10, 5 females and 5 males) and received Nanocovax at single doses of 25 μg, 50 μg, 75 μg and 100 μg or placebo. Untreated mice were used as a control. All animals were regularly monitored continuously within the first 4 hours for behavioural and pathological signs and then daily for the next 14 days for mortality, abnormal behaviour and body weight. At the end of the study, all mice were sacrificed by cervical dislocation and some vital organs, like kidneys, liver and spleen were collected to examine morphological and histological characteristics and weights of the organs.

#### 2.6.2. Repeat-dose toxicity test

According to ICH/GLP guidelines with minor modification, repeat-dose toxicity study was performed on healthy male and female rats with a few modifications. The animals were carefully examined and weighed prior to starting the designed experiment. A total of 36 rats were divided into 6 groups (n = 6; 3 males and 3 females). Rats were intramuscular injected (I.M) Nanocovax daily doses of 25 μg, 50 μg, 75 μg and 100 μg or placebo for 28 days. Untreated rats were used as a control. The mortality and clinical signs was observed daily and the body weight was determined at the indicated time points during the experimental period. The water and food intake was also daily measured. At the end of treatment, all tested rats were anesthetized to collect blood samples for analysis of biochemical and hematological parameters. Following sacrifice of the animals, three vital organs (kidneys, spleen and liver) were immediately isolated, weighed individually and examined histologically.

#### 2.6.3. Local tolerance test

According to ICH/GLP guidelines with minor modification, the local tolerance experiments were performed on New Zealand rabbits. The animals were carefully examined prior to starting the designed experiment. A total of 12 rabbits were divided into 2 groups (n = 6; 3 males and 3 females). The rabbits were intravenously injected (I.V) with a highest dose of Nanocovax (100 μg) and control rabbits were received placebo. Festering and inflammation at the injected position were observed.

### 2.7. Animal ethics statement

This study was carried out in strict adherence to the animal laboratory of Nanogen pharmaceutical Company, the National Institute of drug quality control (NIDQC), and the laboratory of Hanoi Medical University (HMU). The processes were designed according to the guide of ICH/GCP, Drug administration of Vietnam, as well ACTD and approved by Ethics committee of Ministry of Health

### 2.8. Statistical analysis

The collected data were statistically analyzed using Grapthpad Prism, version 5 (Grapthpad Software). Data are expressed as mean ± standard deviation (SD). Statistical analysis was performed using two-way ANOVA analysis with Bonferroni post-tests and one-way ANOVA followed by the Newman-Keuls multiple comparision test to assess the difference between the various groups. Differences described as significant in the text correspond to *p < 0.05; **p < 0.01; ***p < 0.001.

## 3. Results

### 3.1. Recombinant SAR-CoV-2 Spike protein production

To generate SAR-CoV-2 antigen for vaccine development, we designed an optimized DNA sequence encoding the extracellular domain sequence of the Spike protein which has some changes in (1) the S1/S2 Furin cleavage site to minimize the cleavage of S1/S2 during protein production, (2) the two Proline in S2 domain (986-987) to enhance Prefusion-stabilized SARS-CoV-2 Spikes and (3) the 9-arginine residue in the C-terminus (Fig. 1). The recombinant SAR-CoV-2 Spike proteins (S protein)-producing clone was selected on the basis of phenotypic stability, productivity, and key quality properties of the desired product, named CHO-Spike cells. Nucleotide sequence of the gene integrated in the CHO-Spike cell genome was confirmed by sequencing of complementary DNAs (data not shown).

**Figure 1.**
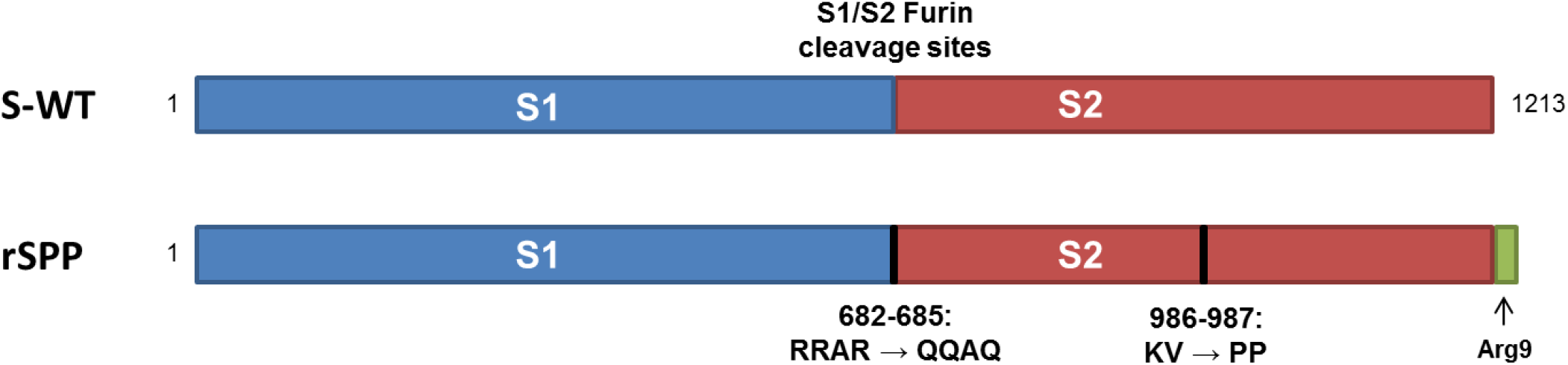
Recombinant SARS-COV-2 Spike Protein Construct: Recombinant S protein (rSPP) construct contain native S protein sequence (aa 1 – 1213) followed by the arginine-9 tag. The S1/S2 furin cleavage sites (RRAR) and 2 amino acid (KV) were mutated as noted.

Upstream production of S protein by CHO-Spike cells was performed in a 500L Bioreactor using a fed-batch process with Feed 3 supplement. The cells were cultured in 380 L PowerCHO-2 medium at initial concentration of 0.5 x 10^6^ cells/mL, and fed with 1.5 % of final volume of Feed 3 daily from day 3 to day 13. The temperature was initiated at 37°C and shifted to 32°C at day 3 of cell culture. Cell density, cell viability and protein titer were determined at indicated time. Figure 2 shows a high performance of CHO-Spike cells in the scale of 500 L using the optimized fed-batch protocol. The CHO-Spike cells could obtain a maximum cell density of 9.48 ± 0.13 ×10^6^ cell/mL (Figure 2A) and survive for 13 days with the cell viability of 80% (Figure 2B). At the end of cultivation, the culture supernatant was harvested and purified for further tests. The data showed that the maximum protein titer was 61.4 ± 1.06 mg/L (Figure 2C) on the harvest day. The SDS-PAGE of harvest sample indicated that the predicted protein was at about 180 kDa, similar with S protein molecular weight (Figure 2D). Besides that, this protein bound specifically to anti-S1 protein antibody (Figure 2E). The harvest sample was purified by conducting on AKTA Pilot 600R. The results of ELISA assay also indicated the purified protein concentration was 21.4 ± 0.18 g per batch with 96.56% of purity (Figure 2H). In addition, residual host cell protein and DNA were not detected in purified S protein (data was not showed).

**Figure 2.**
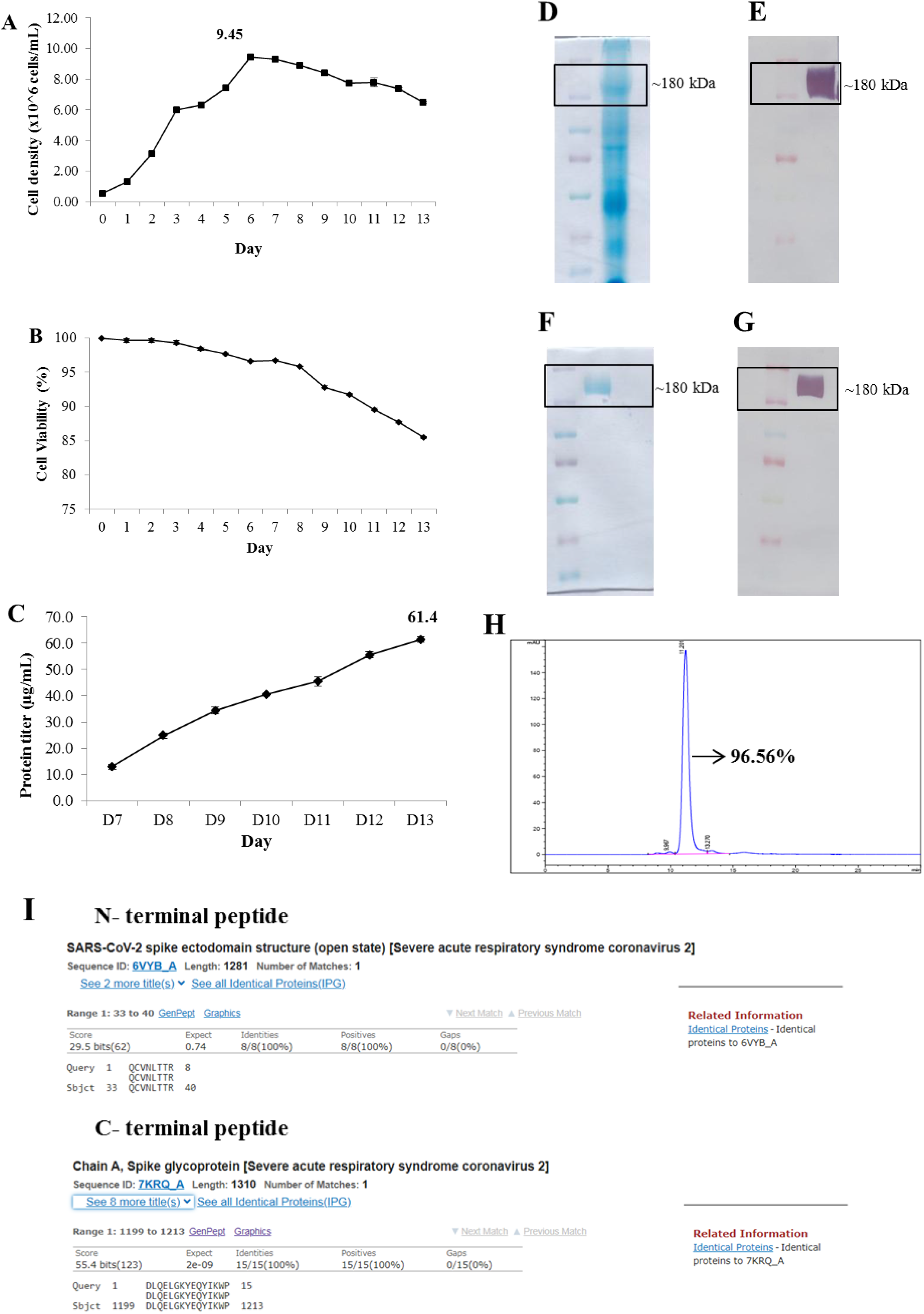
High-yield production and characterization of recombinant SARS-CoV-2 Spike protein: **(A**) Viable cell density (VCD). (**B**) Viability of cells. (**C**) Protein titer from day 7 to end of fed-batch culture. (**D**) SDS-PAGE of harvest sample. (**E**) Western blotting of harvest sample. (**F**) SDS-PAGE of purified S protein. (**G**) Western blotting of purified S protein. (**H**) Purity of S-protein. (**I**) N- and C-terminal peptide sequencing and blasting to complete nonredundant database of protein sequences storage at NCBI

### 3.2. Characterization of recombinant SARS-CoV-2 Spike protein

SDS-PAGE analysis showed that the molecular weight of purified S protein is about 180 kDa, (Figure 2F). Consistent with SDS-PAGE results, anti-S1 antibodies bound specifically to predicted S protein as assayed by Western Blot analysis (Figure 2G). The data of intact mass analysis by MALDI-MS also confirmed that our S protein mass was determined to be 185668 ±1849 Da (Supplementary data S1). On the other hand, the N- and C-terminal sequence and peptide mapping of recombinant S protein were suggested a truncation of N-terminal serine and complete pyroglutamate formation of the N-terminal glutamine. Besides, the C-terminus showed a high heterogeneity with a C-terminal peptide with truncation of AA 1215-1222 as the most abundant variant (Supplementary data S2). Furthermore, the conservation of recombinant S protein was evaluated by comparing N- and C-terminal peptide sequences to complete nonredundant database of protein sequences storage at NCBI, using the BLASTP computer program. The analysis data showed that N- and C-terminal peptide sequences matched 100% to published SARS-CoV-2 Spike protein sequences (Figure 2I).

### 3.3. Nanocovax vaccine formulation

The high purity of S protein-based vaccines requires adjuvant co-administration to enhance immunogenicity [19]. Therefore, SARS-CoV-2 Spike antigen was absorbed into the various concentration of Al^3+^ in Aluminum hydroxide adjuvant. To further evaluate the immunogenicity of Nanocovax, Balb/c mice received intramuscular (IM) injection of PBS (Placebo) or two dose of Nanocovax one week intervally with or without Alhydrogel. On day 14 post-priming injection, blood was collected from the mice to measure humoral immune response by ELISA. As shown in Figure 3, SARS-CoV-2 spike protein-specific IgG titer is highest in mice group receiving 50 μg S protein plus 0.5 mg Al^3+^ (303.29 U/mL) compared with other concentration of Al^3+^, respectively. For further studies, 0.5 mg Al^3+^ was applied to evaluate the immunogenicity of Nanocovax vaccine.

**Figure 3:**
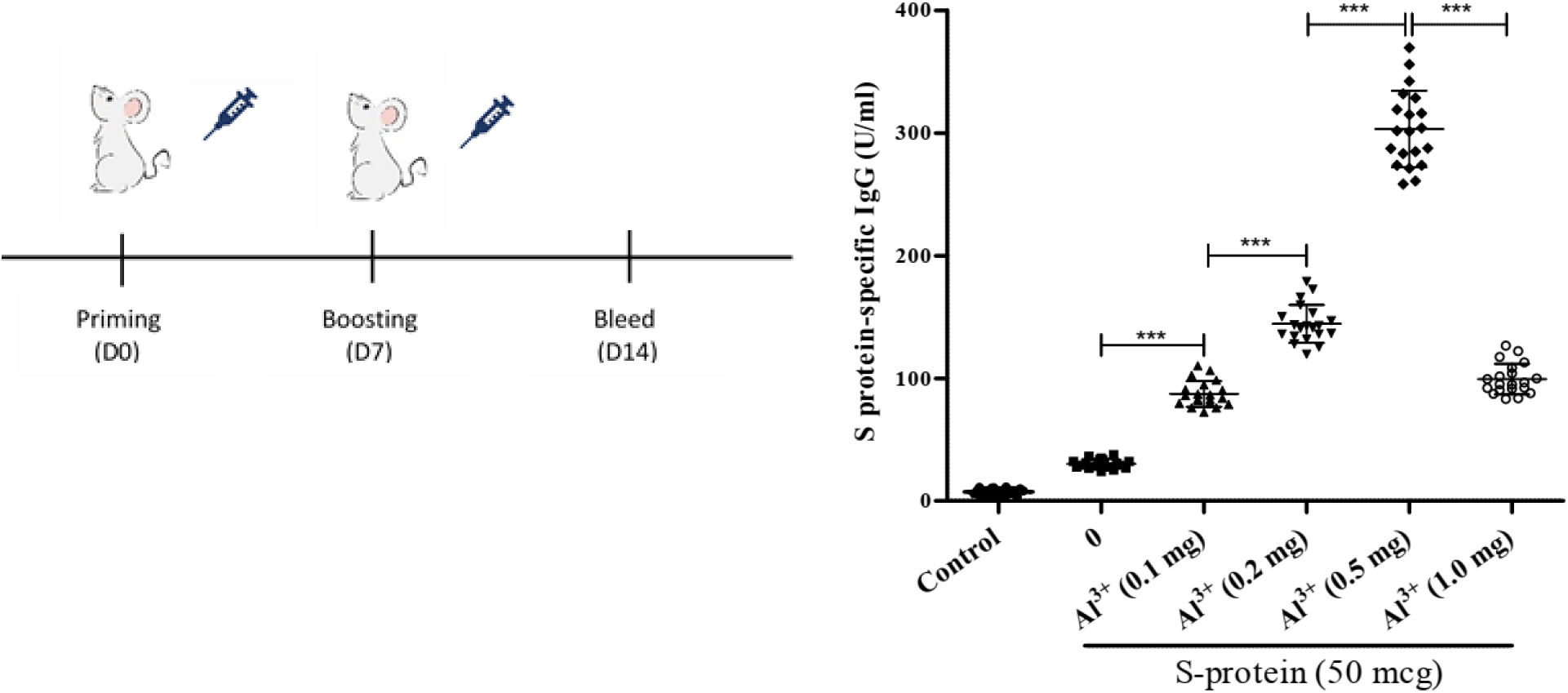
The immunogenicity of Spike protein absorbed into adjuvants using Balb/c mice model (n=20). SARS-CoV-2 Spike protein was absorbed into various concentration of Alhydroxide adjuvant (from 0.1 mg to 1.0 mg Al^3+^). The antibody of IgG levels on sera were analyzed by ELISA on day 14 post-priming. The data represent the mean ± SD and P-values were determined by one-way ANOVA analysis with Newman-Keuls multiple comparision test (***P < 0.001)

### 3.4. Immunogenicity of Nanocovax vaccine in animal models

To assess the immunogenicity of Nanocovax, Balb/c mice were injected with various dosages (25 μg; 50 μg; 75 μg; 100 μg) of vaccine absorbed with 0.5 mg Al^3+^ (aluminum hydroxide adjuvant). The priming and boosting injection were performed on day 0 and day 7, respectively. The blood was collected on day 14 post-priming injection. The levels of total specific IgG were determined by ELISA. The result from fig. 4A shown that IgG levels significantly increased in a dose-dependent manner. To further evaluate the immunogenicity of Nanocovax, Syrian hamsters (*Mesocricetus auratus*) were vaccinated with various doses of Nanocovax (25 μg; 50 μg; 75 μg; 100 μg). The antibodies were detected on day 28 (D28) and day 45 (D45) post-priming. The data from Fig. 4B showed that the amount of S protein-specific IgG in hamster groups that received Nanocovax 25 μg; Nanocovax 50 μg; Nanocovax 75 μg; and Nanocovax 100 μg on D28 were 171.5-fold; 219.8-fold; 222.6-fold; and 253.0-fold compared with the placebo group, respectively. The level of antibodies was slightly decreased on day 45 in the experiments and respective 144.0-fold; 193.8-fold; 240.5-fold; and 196.6-fold compared with the placebo group.

**Figure 4:**
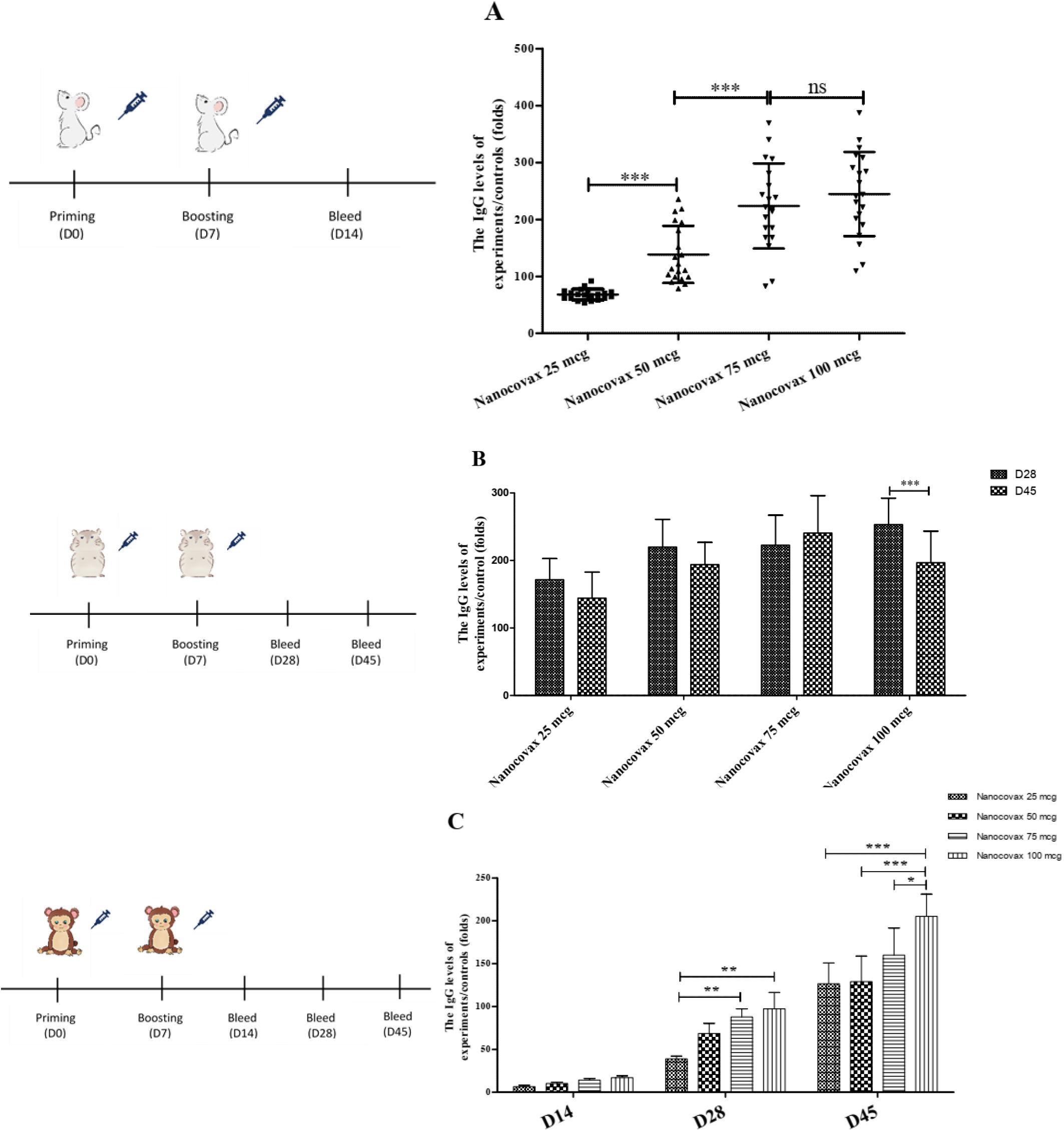
Immunogenicity of Nanocovax vaccine using different animal models. The Balb/c mice (n_group_ = 20) were immunized with various dosages of Nanocovax vaccine (25 μg; 50 μg; 75 μg; and 100 μg). The sera were collected on day 14 and the levels of total specific IgG were determined by ELISA (**A**). Syrian hamsters (n_group_=20) were vaccinated with were immunized with various dosages of Nanocovax vaccine (25 μg; 50 μg; 75 μg; and 100 μg). The sera were collected on D28; D45 and the levels of total specific IgG were determined by ELISA (**B**). The northern pig-tailed macaques (*Macaca leonina*) (n_group_=3) were administrated 02 doses of Nanocovax by I.M injection with 04 dosages: 25 μg; 50 μg; 75 μg; and 100 μg. The sera were collected on day 14; day 28; day 45 and the levels of total specific IgG were determined by ELISA (**C**). The data represent the mean ± SD and P-values were determined by one-way ANOVA analysis with Newman-Keuls multiple comparision test and two-way ANOVA analysis with Bonferroni post-tests (**P < 0.01; ***P < 0.001, ns = not significant)

To confirm the immunogenicity of Nanocovax vaccine, northern pig-tailed macaques (Macaca leonina) were studied. Monkeys were administered 2 doses by I.M injection with various concentrations of Nanocovax (25 μg; 50 μg; 75 μg, 100 μg), or PBS as a negative control. After boosting injection on day 7, the blood was collected on day 14; day 28, and day 45 to detect the level of antibodies. The results are illustrated in fig 4C. These data indicated that the levels of IgG in monkeys in the groups that received Nanocovax 25 μg; Nanocovax 50 μg; Nanocovax 75 μg and Nanocovax 100 μg were significantly increased in a time-dependent manner. On day 28 post-priming injection, the IgG levels were slightly increased in all groups that received Nanocovax vaccines and were 39.02-fold; 68.58-fold; 87.82-fold; and 97.37-fold, respectively. Similar, on day 45, the amount of IgG in the serum of monkeys in all vaccinated groups was extremely significantly increased compared with the control group (The IgG levels of vaccinated animals were 126.5-fold; 129.1-fold; 159.95-fold; and 205.12-fold, respectively).

On the other hand, sera were collected from vaccinated animal models and the percent of neutralizing antibodies was measured by a surrogate virus neutralization test (SARS-CoV-2 sVNT Kit). For the Balb/C mice model, mice were immunized with 2 doses of vaccine, 7 days apart. Day 14 post-priming injection, blood was collected from the mice to detect the percent of inhibition. As shown in Fig. 5A, four dosages of Nanocovax (25 μg; 50 μg; 75 μg; and 100 μg) strongly inhibited SARS-CoV-2 virus compared with PBS group (***P < 0.001). The inhibition by Nanocovax 50 μg was not significantly different from Nanocovax 75 μg (89.60% versus 89.40%). A significant difference was observed between the 25 μg group compared with the 50 μg group (77.5% versus 89.60%), as well as the 75 μg group versus the 100 μg group (89.40% versus 97.30%). For the Syrian Hamster model, blood in hamsters was collected on day 28 and day 45 post-priming injection with four dosages of Nanocovax including 25 μg; 50 μg; 75 μg; and 100 μg. The percent of inhibition was detected by SARS-CoV-2 sVNT Kit. The results from Fig. 5B indicated that no significant difference (P > 0.05) was observed among all groups on day 28 and day 45 post-priming injection with four dosages of Nanocovax vaccine (25 μg; 50 μg; 75 μg; and 100 μg) respectively. Otherwise, vaccinated hamster groups showed significant inhibition by antibodies in sera compared with the non-vaccinated hamster group (***P<0.001). We next measured the virus neutralization ability of antibodies in the non-human primate model. The monkeys (Macaca leonina) were immunized twice a week with Nanocovax with dosages of 25 μg, 50 μg, 75 μg, and 100 μg. Sera were collected on day 14, day 28, and day 45 after priming I.M injection to determine inhibition of S protein using the virus surrogate neutralization KIT. The results from fig. 5C show that the monkeys vaccinated with the 25 μg dosage of Nanocovax produced no significant inhibition on day 14. (The percent of inhibition was 26.14%, lower than the 30% Cutoff value). On day 28 and day 45, the monkey groups immunized with four dosages of Nanocovax had positive results (> 30% Cutoff value) and no significant difference in the percent of inhibition.

**Figure 5:**
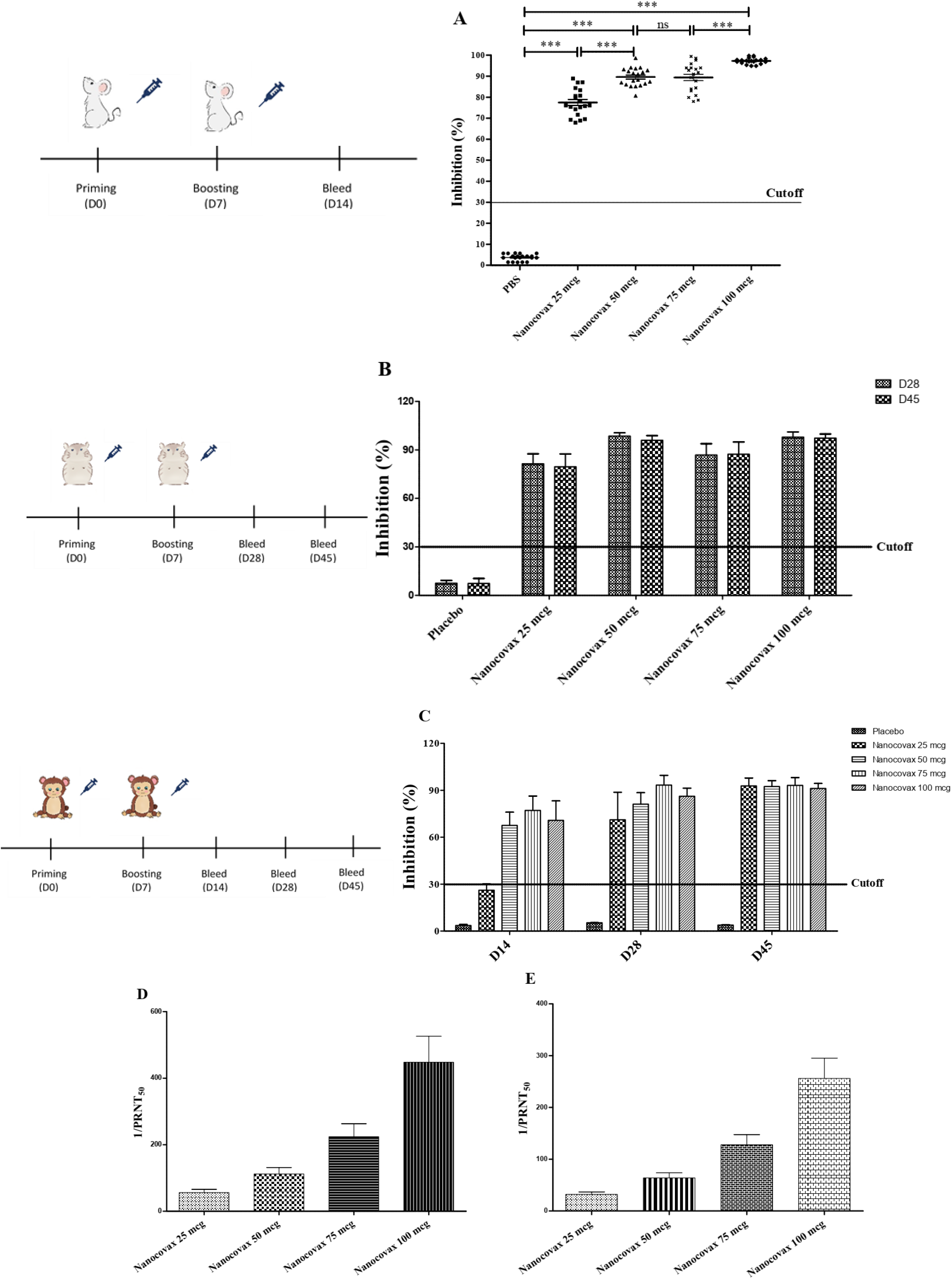
The SARS-CoV-2-neutralizing antibodies: The Balb/c mice (n_group_ = 20) were immunized with various dosages of Nanocovax vaccine (25 μg; 50 μg; 75 μg; and 100 μg) and the sera were collected on day 14. The levels of total specific IgG were determined by ELISA **(A)**. The SARS-CoV-2-neutralizing antibodies in sera were measured by PRNT_50_ (**D**). Syrian hamsters (n_group_=20) were vaccinated with were immunized with various dosages of Nanocovax vaccine (25 μg; 50 μg; 75 μg; and 100 μg). The sera were collected on D28; D45 and the levels of total specific IgG were determined by ELISA (**B**). The SARS-CoV-2-neutralizing antibodies in sera were measured by PRNT_50_ (**E**). The northern pig-tailed macaques (*Macaca leonina*) (n_group_=3) were administrated 02 doses of Nanocovax by I.M injection with 04 dosages: 25 μg; 50 μg; 75 μg; and 100 μg. The sera were collected on day 14; day 28; day 45 and the levels of total specific IgG were determined by ELISA (**C**). The data represent the mean ± SD and P-values were determined by one-way ANOVA analysis with Newman-Keuls multiple comparision test and two-way ANOVA analysis with Bonferroni post-tests (**P < 0.01; ***P < 0.001, ns = not significant)

Furthermore, SARS-CoV-2-neutralizing antibodies can be measured by the PRNT_50_ viral neutralization assay, which is used to determine the levels of neutralizing antibodies in sera from the vaccinated animals. The results from fig. 5D showed that the specimens of Balb/C immunized with Nanocovax 25 μg; 50 μg; 75 μg; and 100 μg had respective SARS-CoV-2-neutralizing antibodies (1/PRNT_50_) titer from 40 to 640. Similarly, all specimens of hamsters received Nanocovax 25 μg; 50 μg; 75 μg; and 100 μg had respective SARS-CoV-2-neutralizing (1/PRNT_50_) titer from 20 to 320 (Fig. 5E).

### 3.5. Protective efficacy of Nanocovax vaccine in hamster models

All procedures were performed in BSL-3 facility at NIHE (National Institute of Hygiene and Epidemiology) Vietnam. The hamsters were assigned to the following groups: (1) vaccinated with Nanocovax on day 0 and day 7 and then challenged with the high level of the SARS-CoV-2 virus on day 14 by the intranasal route (TCID_50_ = 2×10^5^); (2) Placebo injection with PBS and challenged with the high level of the SARS-CoV-2 virus on day 14 by the intranasal route (TCID_50_ = 2×10^5^); (3) vaccinated with Nanocovax on day 0 and day 7 and then challenged with the low level of the SARS-CoV-2 virus on day 14 by the intranasal route (TCID_50_ = 1×10^3^), and (4) placebo injection with PBS and challenged with the low level of the SARS-CoV-2 virus on day 14 by the intranasal route (TCID_50_ = 1×10^3^). Animals were monitored for signs of morbidity (such as weight loss, ruffled hair, sweating, etc.) for 14 days after challenge with live SARS-CoV-2. The observation from fig. 6A, 6B shown that three vaccinated hamsters from each group had no signs of weight loss: weight maintenance for 1-2 days post-infection and weight gain continued from day 3 post-infection and day 14 post-infection (11.8% to 14.5%). No symptoms were observed in vaccinated animals: no shortness of breath, ruffled fur, or lethargy were observed in any vaccinated hamsters receiving the low and high doses of SARS-CoV-2.

**Figure 6:**
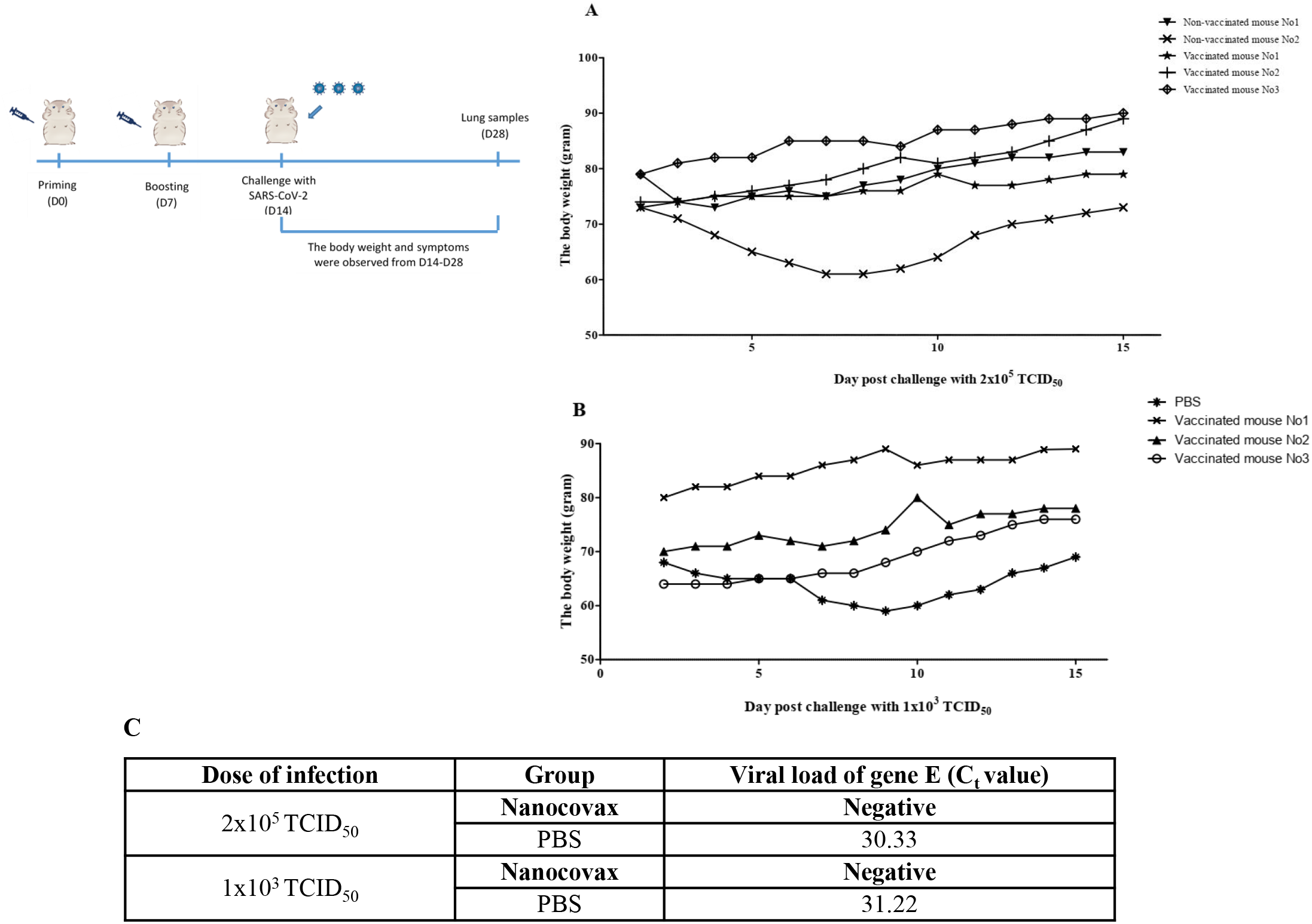
Challenge of Syrian hamsters (*M. auratus*) with live SARS-CoV-2. After vaccination, the hamsters were challenged with low levels and high levels of SARS-CoV-2 virus. The weight gain in hamster was monitored after challenge with high levels (**A**) and low levels of SARS-CoV-2 (**B**). SARS-CoV-2 load (gene E) in lung samples 14 days post-challenge was determined by Real-time RT-PCR (**C**).

Three control hamsters exhibited ruffled fur, lethargy, and sweating symptoms on day 1 and 2 after the virus challenge. Two of three animals showed severe weight loss by day 7 or 8 post-infection (13.2% to 16.4%), and gained weight slowly from day 8 or 9 after the challenge test. On day 28, the lungs from the vaccinated hamster group and the non-vaccinated group were collected to detect SARS-CoV-2 by Realtime RT-PCR. The results in fig. 6C showed that no SARS-CoV-2 virus-specific RNA was detected in lung samples of the vaccinated group after 14 days of challenge with viral concentrations 2×10^5^ TCID_50_ and 1×10^3^ TCID_50_ compared with the non-vaccinated group (Ct = 30.33, and Ct = 31.22, respectively).

### 3.6. Safety evaluation of Nanocovax vaccine

#### 3.6.1. Single-dose toxicity

In the single-dose toxicity test, the mice were injected intramuscularly with Nanocovax at dosages of 25 μg, 50 μg, 75 μg and 100 μg and were observed up to 14 days. Control mice received placebo. The result of the study showed no mortality and no drug-related toxicity signs in the tested mice at all tested doses within 14 days. Moreover, there were no significant differences between groups for body weights and relative organ weights (Fig. 7A; 7B). Macroscopic examination showed that there were no abnormalities in physical appearance of liver, heart, kidneys, spleen, lungs and intestines between the treated groups and the control group (Fig. 7C). Histological analysis shows no sign of congestion or necrosis in the liver, kidneys, and spleen (Fig. 7D).

**Figure 7:**
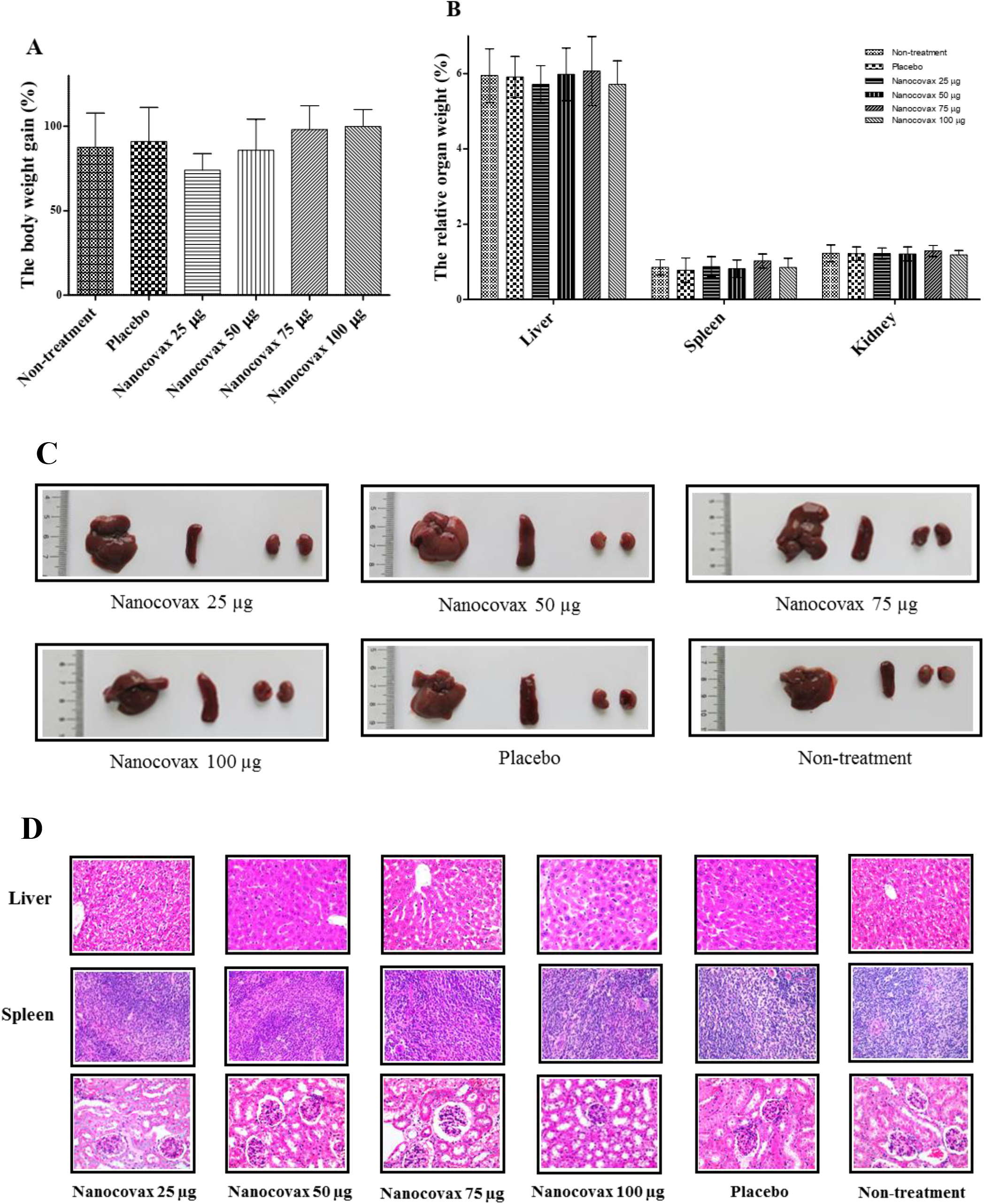
The body weight gain (**A**) and the relative organs weight (**B**). Anatomical structure of organs (**C**) and histological analysis of liver, spleen, kidney in single-dose test (**D**). The data represent the mean ± SD and P-values were determined by one-way ANOVA analysis with Newman-Keuls multiple comparision test, and two-way ANOVA analysis with Bonferroni post-tests (ns = not significant).

#### 3.6.2. Repeat-dose toxicology

In the repeat-dose study, the male and female rats were injected intramuscularly once a day for 28 consecutive days with with Nanocovax at dosages of 25 μg, 50 μg, 75 μg and 100 μg. Control rats received placebo. The treated rats and the controls appeared uniformly healthy and no lethality was recorded in tested rats during the 28-day treatment period. There were no clinically abnormal symptoms in general behaviour between treatment and control groups. In comparison with control rats, the body weight of female and male rats gradually increased during the test period. However, there was no statistically significant difference in body weight gain between the treated groups and the control group (Fig. 8A). Similarly, no significant difference was observed in organ weights of rats, both males and females between treated groups and control group (Fig. 8B). The results showed that during the treatment, vital organs such as the liver and kidney were not adversely affected. Macroscopic observations showed that there were no lesions and no abnormalities of physical appearance of liver, kidneys and spleen observed in all groups. As illustrated in figure 8C, no significant macroscopic changes, such as colour, size, shape and texture, in liver, spleen, and kidneys were observed between groups. The indicators of the hematological parameters and biochemical parameters of rats are shown in Fig 9A; 9B, respectively that had no significant difference in vaccinated groups and control groups. Similarly, urinalysis of rats also had no significant difference in vaccinated groups and control groups (data not shown). The rerults from histopathological analysis revealed that none of kidney, liver and spleen tissues of both male and female rats that received Nanocovax daily at all doses showed any damage or pathological alterations under microscopic observation (Figure 9C).

**Figure 8:**
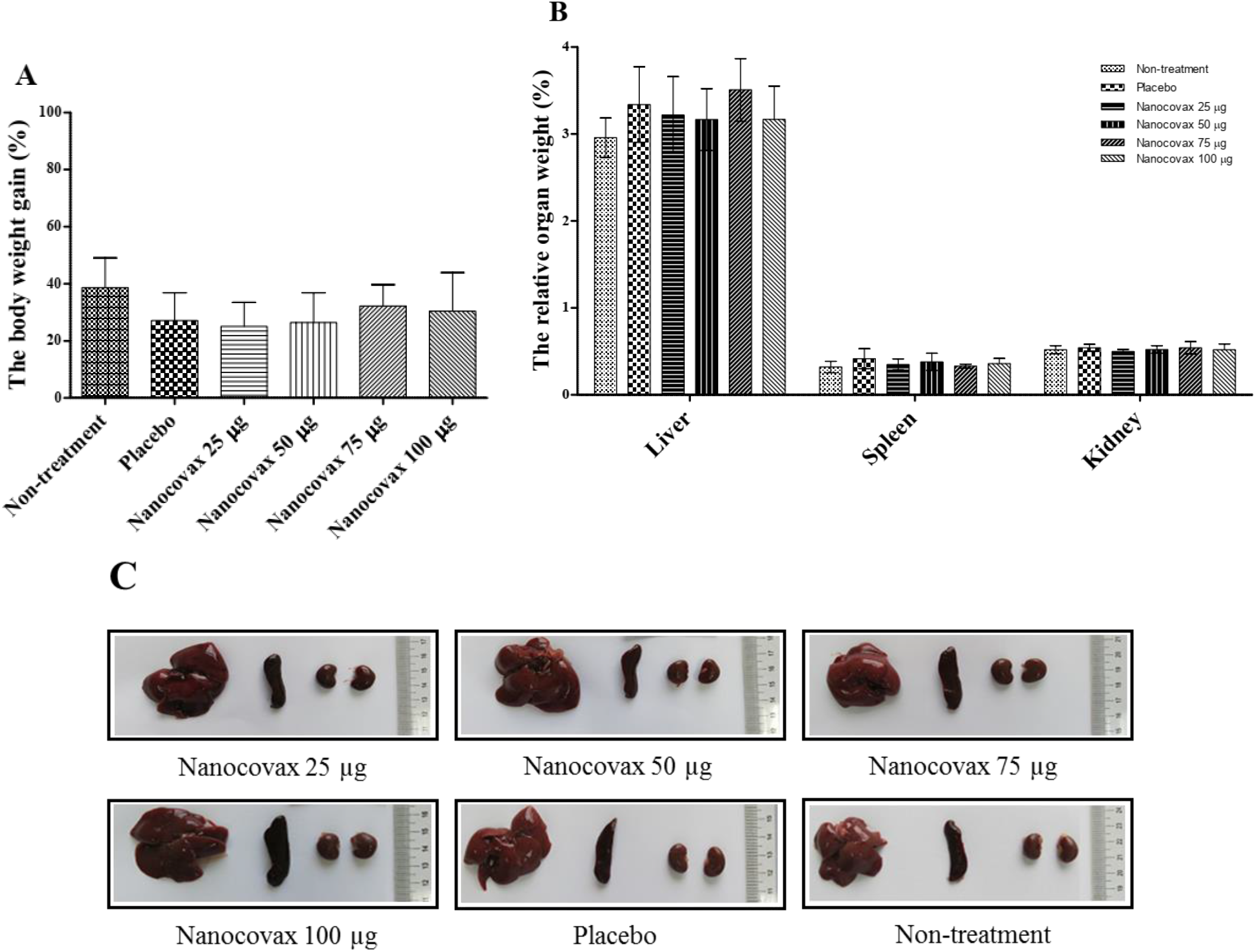
The body weight gain (**A**), the relative organs weight (**B**) and anatomical structure of organs of rat (*Rattus norvegicus*) (**C**) in repeat-dose toxicology. The data represent the mean ± SD and P-values were determined by two-way ANOVA analysis with Bonferroni post-tests, one-way ANOVA analysis with Newman-Keuls multiple comparision test

**Figure 9:**
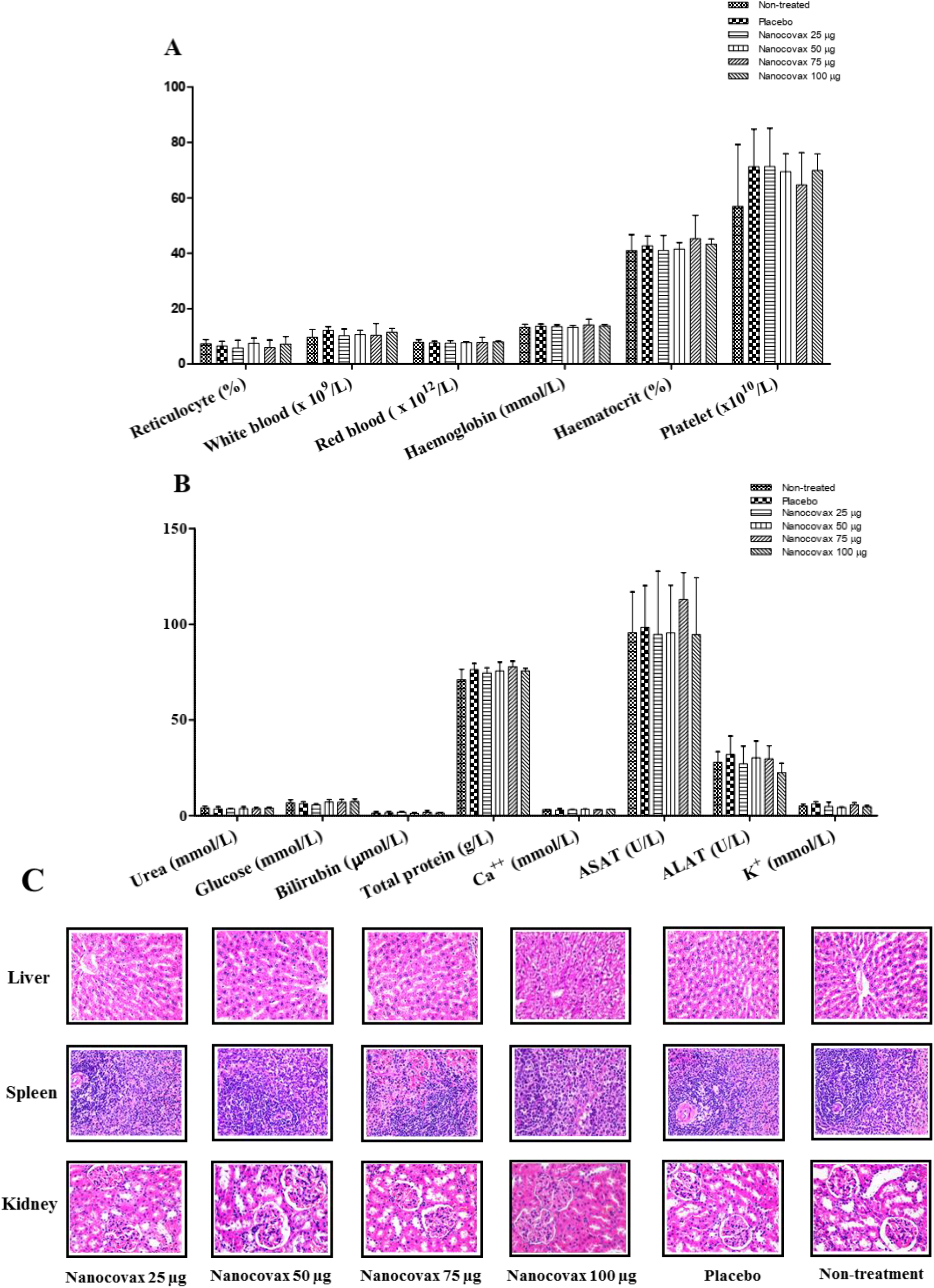
The indicators of the hematological parameters (**A**) and biochemical parameters (**B**). Histological analysis of liver, spleen, kidney in repeat-dose test (magnification 40X) (**C**). The data represent the mean ± SD and P-values were determined by two-way ANOVA analysis with Bonferroni post-tests.

#### 3.6.3. Local tolerance

The experiments were performed on New Zealand rabbits. The rabbits (n=6) were intravenously injected (I.V) with a sigle-dose of Nanocovax 75 μg and other rabbits group (n=6) were received PBS as a placebo. Before injection, rabbit ears were normal, and the injection sites, body temperatures, and bodyweights of all animals were monitored for 7 days. 2 hours after injection, 1 of 6 rabbits in each of the experiment groups was slightly swollen and light red at the injection site. 4 hours later, 3/6 rabbits in the experiment group and 4/6 rabbits in the placebo group were slightly swollen and light red at the injection site. After 24 hours, 2/6 rabbits in the experiment group were slightly swollen and light red at the injection site; 3/6 rabbits in the experiment group were swollen and red at the injection site. For the placebo group, 1/6 rabbits was slightly swollen and light red; 5/6 rabbits were swollen and red at the injected site. After 48 hours, 3/6 rabbits of the experiment group were swollen and red compared with the placebo group in which 4/6 rabbits were slightly swollen and light red and 2/6 rabbits were swollen and red at site of injection. 72 hours later, 3/6 rabbits were slightly swollen and light red at the site of injection in each group. 96 hours later, only one rabbit was slightly swollen and light red at the site of injection in the two groups. After 144 hours, all adverse events were fully recovered at the site of injection on the ear (data not shown). Furthermore, the bodyweights of rabbits were monitored on day −1, day 1, day 4, and day 7 after challenging with high dose of Nanocovax as well as PBS. The result from fig. 10 showed that the bodyweight of rabbits in all experiments slightly increased but the increase of bodyweight before versus after injection was not significantly different and there was no significant difference among all animal groups.

**Figure 10:**
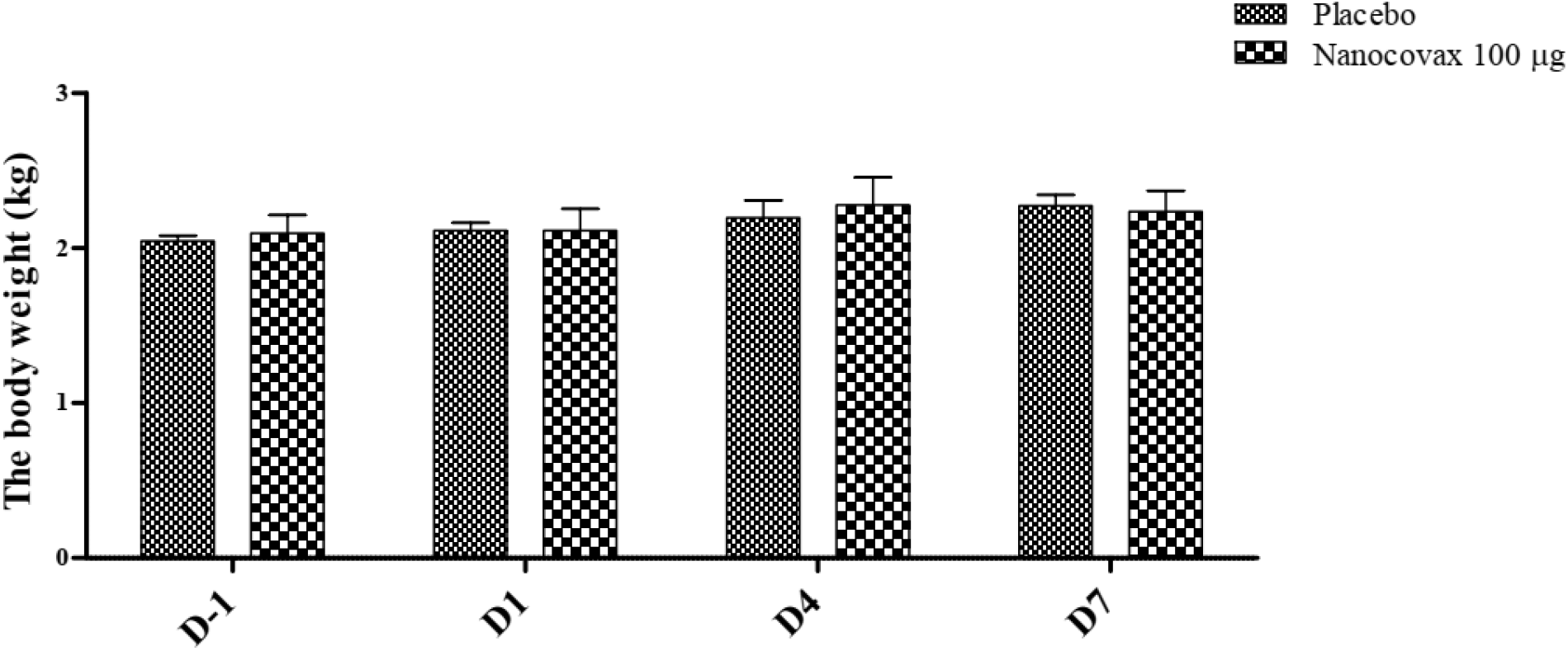
The body weight gain of New Zealand rabbits in local tolerance. The data represent the mean ± SD and P-values were determined by two-way ANOVA analysis with Bonferroni post-tests.

## 4. Discussion

The development of vaccines with high immunogenicity and safety are the major concerns for control of the global COVID-19 pandemic and protection of further illness and fatalities. Vaccine candidates are currently under development using different platforms, such as inactivated vaccines, live-attenuated vaccines, viral vector (adenovirus) vaccines, DNA vaccines, and mRNA vaccines. In other platforms, recombinant protein vaccines were used together with adjuvants to enhance adaptive immunity. Aluminum salt-based adjuvants are approved for many human vaccines such as Hepatitis A; Hepatitis B; Haemophilus influenza type b (Hib); Diphtheria-tetanus-pertusis (DtaP, Tdap) [20]. Currently, Aluminum adjuvants have been applied in SARS-CoV-2 vaccines such as RBD vaccine; spike protein vaccine; inactivated virus vaccine; VLP vaccine [21]. For Nanocovax vaccine, spike protein was designed (Fig. 1) and firstly harvested at final concentration (21.06 g per batch) after purificattion with 96.56% purity (Fig. 2). Secondly, spike protein was formulated with aluminum hydroxide adjuvant for pre-clinical research. Here, the results from fig. 3 showed that spike protein absorbed on aluminum hydroxide adjuvant-induced significantly higher IgG levels compared to mice group vaccinated with spike protein only, as well control group. Based on these results, aluminum hydroxide adjuvant was suggested for further experiments.

To evaluate the immunogenicity of Nanocovax vaccine, three animal models including Balb/C mice; Syrian hamster mice, and non-human primates were used. The antibody IgG titer in sera from vaccinated Balb/C groups with various concentrations of Nanocovax (25 μg; 50 μg; 75 μg; 100 μg) significantly increased in a dose-dependent manner on D14 (fig. 4A). However, the levels of IgG titer were similar on D28, as well D45 by using the hamsters model (fig. 4B). In addition, the amount of IgG in serum of monkeys in all vaccinated groups was significantly in a time-dependent manner (fig. 4C).

The neutralizing antibodies to SARS-CoV-2 are urgently needed to determine not only the infection rate, immunity, but also vaccine efficacy during clinical trials [15]. Sera were collected from vaccinated animal models, as well non-human primates to measured the percent of neutralizing antibodies by surrogate virus neutralization test. The results from fig 5 showed that the antibody in sera can neutralize SARS-CoV-2 virus with a high percent of inhibition. Agreement with result from Bošnjak [22], a strong positive correlation between surrogate virus neutralization test and the levels of S-specific IgG. Our animal models showed that the IgG levels in sera from vaccinated animals with Nanocovax increased high affinity viral neutralizing antibodies. Futhermore, SARS-CoV-2-neutralizing antibodies can be measure by PRNT_50_. Here, sera from Balb/C and Hamster models had neutralizing antibodies titer from 80 to 640, and 20 from 320, respectively.

Next, the protective efficacy of Nanocovax vaccine on hamster was used in study. After challenging with SARS-CoV-2 virus, no symptoms were observed such as shortness of breath, ruffled fur, or lethargy as well as weight loss in all vaccinated mice verse control hamster groups. On day 28, SARS-CoV-2 virus-specific RNA was detected in the lung samples of non-vaccinated groups by Real-time RT-PCR (fig. 6)

The safety of the Nanocovax vaccine was investigated based on studies of single-dose toxicity; repeat-dose toxicity; as well local tolerance test. The results from fig. 7 and fig. 8 showed that Nanocovax has no single-dose toxicity effect testing on *Mus musculus* var. Albino mice. Similarly, the results from fig. 9; fig. 10 showed that Nanocovax vaccine has no repeated-dose toxicity effect testing on Rat (*Rattus norvegicus*) with three dosages (25 μg; 50 μg; 75 μg). Furthermore, Nanocovax also produced no local irritation in New Zealand rabbits at the highest dose of 100 μg (fig. 10).

Taken all together, Nanocovax vaccine demonstrated immunogenicity and safety based on animal models as well as non-human primate model. These results support the clinical phase I and phase II development of the Nanocovax vaccine.

## Abbreviations

COVID-19: Coronavirus disease-2019
SARS-CoV-2: Severe Acute Respiratory Syndrome coronavirus 2
RBD: receptor-binding domain
ACE-2: angiotensin-converting enzyme 2
CHO: Chinese hamster ovary

## Acknowledgments

This study was supported by Ministry of Science and Technology, Vietnam under grant number: ĐTĐL.CN.51/20.

## Author contributions

The authors contributed equally to this work

